# An Optimized Registration Workflow and Standard Geometric Space for Small Animal Brain Imaging

**DOI:** 10.1101/619650

**Authors:** Horea-Ioan Ioanas, Markus Marks, Valerio Zerbi, Mehmet Fatih Yanik, Markus Rudin

## Abstract

The reliability of scientific results critically depends on reproducible and transparent data processing. Cross-subject and cross-study comparability of imaging data in general, and magnetic resonance imaging (MRI) data in particular, is contingent on the quality of registration to a standard reference space. In small animal MRI this is not adequately provided by currently used processing workflows, which utilize high-level scripts optimized for human data, and adapt animal data to fit the scripts, rather than vice-versa. In this fully reproducible article we showcase a generic workflow optimized for the mouse brain, alongside a standard reference space suited to harmonize data between analysis and operation. We introduce four separate metrics for automated quality control (QC), and a visualization method to aid operator inspection. Benchmarking this workflow against common legacy practices reveals that it performs more consistently, better preserves variance across subjects while minimizing variance across sessions, and improves both volume and smoothness conservation RMSE approximately 2-fold. We propose this open source workflow and the QC metrics as a new standard for small animal MRI registration, ensuring workflow robustness, data comparability, and region assignment validity, all of which are indispensable prerequisites for the comparability of scientific results across experiments and centers.

## Background

Correspondence of brain areas across individuals is a prerequisite of generalizable statements regarding brain function and organization. This is achieved by spatial transformation of brain maps in a study to a population or study reference template. This process, called registration, is integral to any neuroimaging workflow attempting to produce results which are both spatially resolved and meaningful at the population level.

The computations required for registration are com-monly performed in the preprocessing workflow, though image manipulation may only take place once inter-subject comparison becomes needed. As a consequence of this peripheral positioning in the data evaluation sequence, and of the independence from experimental designs and hypotheses, registration is often relegated to default values and exempt from rigorous design efforts and QC.

Registration in human brain imaging benefits from high-level functions (e.g. `flirt` and `fnirt` from the FSL package[1], or `antsIntroduction.sh` from the ANTs package[2]), optimized for the size and spatial features of the human brain. In small animal brain imaging, registration is performed via the selfsame high-level functions as human brain imaging — rendered usable by adjusting the data to fit function parameters, rather than vice-versa. This approach compromises data veracity, limits optimization potential, and represents a notable hurdle for the methodological improvement of small animal brain imaging.

Ad-hoc workflows for preclinical imaging often emerge as the need mandates, but lack broad reusability due to highly variable protocols for data acquisition across centers [3]. While further preprocessing advancements are being made for general-purpose human data preprocessing [4], the preclinical field still lacks similarly reliable tools. Recently, a preclinical registration framework has been released [5], dealing specifically with structural recordings focused on voxel-based morphometry (VBM) and validated via phantom-based metrics. General-purpose registration workflows applicable both to functional and structural data remain lacking, as do benchmarking efforts which take into account QC metric challenges arising from the heterogeneity of in vivo experiments.

Below, we explicitly describe current practices used in extant ad-hoc workflows, in an effort to not only propose better solutions, but do so in a falsifiable manner which provides adequate detail for both novel and the legacy methods.

### Manipulations

The foremost data manipulation procedure in present-day small animal MRI is scale adjustment. Scale is represented by the Neuroimaging Informatics Technology Initiative format (NIfTI) [6] affine transformation parameters, which map the data matrix to spatial coordinates. Commonly, this manipulation constitutes a 10-fold increase in each spatial dimension.

In order to produce acceptable results from brain extraction based on human priors, it may be necessary to additionally adjust the data matrix content itself. This may involve applying an ad-hoc intensity-based percentile threshold to delete non-brain as well as anterior and posterior brain voxels, and leave a more spherical brain for human masking functions to operate on. While conceptually superior solutions adapting parameters to animal data are available [7, 8] and might remove the need for this step of data adaptation, rudimentary solutions remain popular. Both these function adaptations for animal data and the animal data matrix content adaptations for use with human brain extraction functions are, however, prone to completely or partially remove the olfactory bulbs. For this reason, the choice is sometimes made to simply forego brain extraction.

Scan orientation metadata may be seen as problematic, and consequently deleted. This consists in resetting the NIfTI S-Form affine to zeroes, and is intended to mitigate orientations incompatible with the target template. While it is true that scanner-reported affine spaces for small animal data may be nonstandard (the confusion is two-fold: small animals lie prone with the coronal plane progressing axially whereas higher primates lie supine with the horizontal plane progressing axially), affine spaces of small animal brain templates may be nonstandard as well. A related manipulation is dimension swapping, which changes the order of the NIfTI data matrix dimensions rather than the affine metadata. Thus, correct or automatically redressable affine parameters can be deleted and data degraded beyond easy recovery, in order to correspond to a malformed template.

### Templates

As the above demonstrates, the template is a key dependency for registration. Templates used for small animal brain MRI registration are heterogeneous and include histological, as well as ex vivo MRI templates, scanned either inside the intact skull or after physical brain extraction.

Histological templates benefit from high spatial resolution and, in the mouse model in particular, access to molecular information in the same coordinate space. Such templates are not produced in volumetric sampling analogous to MRI, are hard to register to due to their contrast, may be severely deformed due to their extraction process, and are often not assigned a meaningful affine transformation after conversion to NIfTI. Data registered to templates thus deformed is challenging to use for navigating the intact brain during stereotactic surgery.

Ex vivo templates based on extracted brains share most of the deformation issues present in histological templates; they are, however, available in MR contrasts, making registration more reliable. Ex vivo templates based on intact heads provide both MR contrast and brains largely free of deformation and supporting whole brain registration.

### Challenges

The foremost challenges in small animal MRI registration consist in eliminating data-degrading workarounds, reducing reliance on high-level interfaces with inappropriate optimizations, and improving the quality of standard space templates. Information loss (pertaining to both the affine and the data matrix) is a particularly besetting issue, since the loss of data at the onset of a neouroimaging workflow persists throughout downstream steps and precludes numerous modes of analysis (fig. 1a vs. fig. 1b). Additionally, given data heterogeneity in various MRI applications, processing effects need to be evaluated with respect to at least the most common contrasts, such as anatomical *T*_1_ and *T*_2_-weighted imaging, as well as variations of functional contrast, such as cerebral blood volume (CBV) and blood-oxygen-level-dependent (BOLD) imaging.

**Figure 1:**
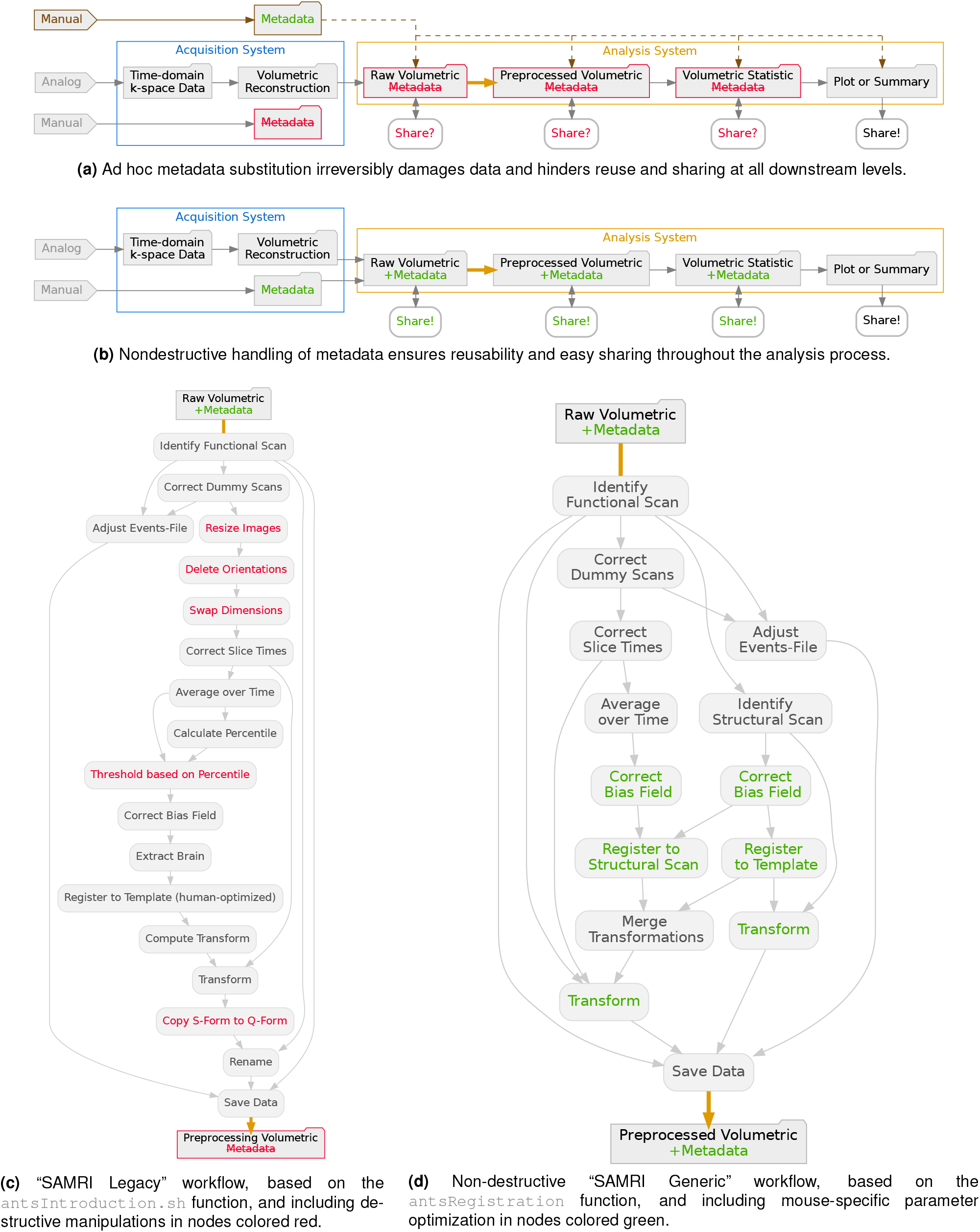
The SAMRI Generic workflow uses fine-tuned animal priors to enhance registration quality while preserving metadata integrity. Directed graphs depict both the overall context of MRI data processing and analysis (**a**,**b**), as well as the internal structure of the two registration workflows compared in this article (**c**,**d**), which insert into the broader context at the bold orange arrow positions. Technical detail available in fig. S5.

## The Optimized Workflow

Processing workflow complexity should be manageable to prospective users with only cursory programming experience. However, workflow transparency, sustainability, and reproducibility should not be compromised for trivial ancillary features. We thus abide by the following design guidelines: (1) *each workflow is represented by a high-level function*, with operator-understandable parameters, detailing operations performed, rather than implementations; (2) *workflow functions are highly parameterized but include workable defaults*, so users may significantly adapt workflows without editing constituent code; (3) *graphical or interactive interfaces are avoided*, as they impede reproducibility, encumber the dependency graph, and reduce project sustainability.

Given the aforementioned principles, we have constructed two registration workflows: The “Legacy” workflow (fig. 1c, exhibiting the common practices detailed in the Background section) and the novel “Generic” workflow (fig. 1d), Both workflows start by performing dummy scan correction on the fMRI data and the stimulation events file, based on BIDS [9] metadata, if available. The “Legacy” workflow subsequently applies a 10-fold multiplication to the voxel size, and deletes the orientation information from the affine matrix. Further, dimensions are swapped so that the data matrix matches the RPS (left *→* Right, anterior *→* Posterior, inferior *→* Superior) orientation of the corresponding template (fig. 2b). Following these data manipulation steps, a temporal mean is computed, and an empirically determined threshold (10 % of the 98^th^ percentile) is applied. Subsequently, the bias field is corrected and masked using the `fast` and `bet` FSL functions. The image is then warped into the template space using the `antsIntroduction.sh` ANTs function. Lastly, affine variants are harmonized. The “Generic” workflow follows up on dummy scan correction with slice timing correction, computes the temporal mean of the functional scan, and bias corrects the temporal mean — using the `N4BiasFieldCorrection` ANTs function, with spatial parameters adapted to the mouse brain. Analogous operations are performed on the structural scan, following which the structural scan is registered to the reference template, and the functional scan temporal mean is registered to the structural scan — using the `antsRegistration` ANTs function, with spatial parameters adapted to the mouse brain. The structural-to-template and functional-to-structural transformation matrices are merged and applied in one warp step to the functional data, while the structural data is warped solely via the structural-to-template transformation matrix.

**Figure 2:**
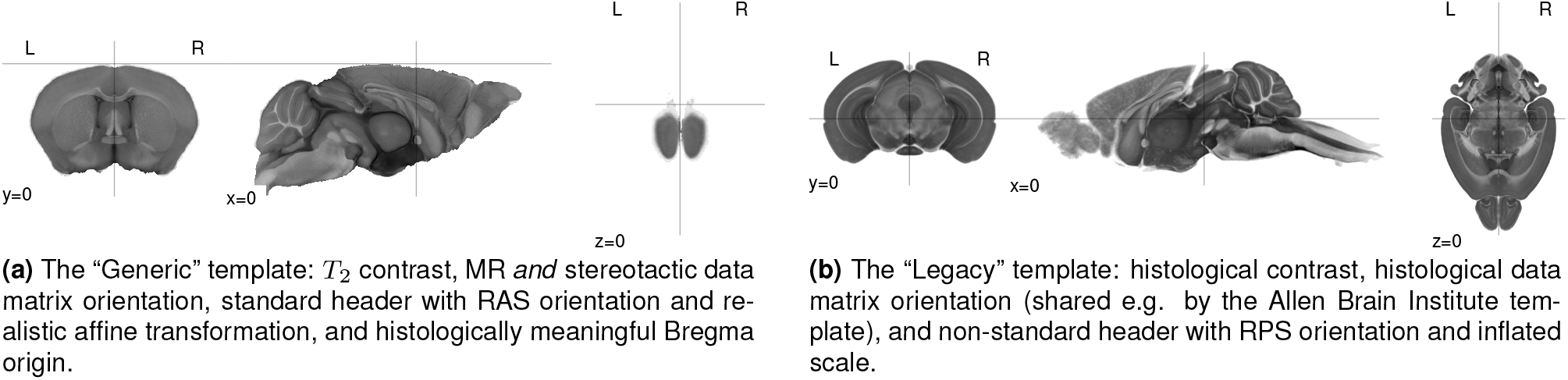
The “Generic” template provides canonical orientation and Bregma centering. Illustrated are multiplanar depcitions of the “Generic” and “Legacy” mouse brain templates, with slice coordinates centered at zero on all axes.

The workflow is implemented in Python, with automatically generated Command Line Interfaces (CLIs) available for Bash. As registration is indispensable to data analysis (rather than a stand-alone process), the workflows are distributed as part of a comprehensive workflow package, SAMRI [10].

### Template Package

Target template quality is highly consequential to registration performance and suitability as a standard procedure. For example, an inflated template size mandates according parameters for all functions handling data in affine space. Further, if template axes are flipped, the first (rigid) registration step may be incorrectly determined and the image deformed to match the template at an incorrect orientation. Consequently, template quality must be ascertained, and a workflow-compliant default should be provided.

Our recommended template (fig. 2a) is derived from the DSURQE template of the Toronto Hospital for Sick Children Mouse Imaging Center [11]. We shift its geometric origin to the Bregma landmark, providing harmonization with histological atlases and surgical procedures. The template is in the canonical NIfTI orientation, RAS (left *→* Right, posterior *→* Anterior, inferior *→* Superior), and has a coronal slice positioning typical for both MRI and stereotactic surgery. All of the associated files are identified with the prefix `dsurqec` in the template package.

We bundle the aforementioned MR template with histological templates, derived from the Australian Mouse Brain Mapping Consortium (AMBMC) [12] and Allen Brain Institute (ABI) [13] templates, which provide ample rostrocaudal coverage and serve as a reference for numerous gene expression and projection maps, respectively. We reorient the AMBMC template from its original RPS orientation to the canonical RAS, and apply an RAS orientation to the orientation-less ABI template after NIfTI conversion from the original NRRD format. Corresponding files are prefixed with `ambmc` and `abi`, respectively.

Additionally, we provide DSURQEC and AMBMC derived templates, in the widespread but incorrect RPS orientation, with the aforementioned tenfold increase in voxel size. These are prefixed with `ldsurque` and `lambmc`, respectively.

All templates are provided at 40 µm and 200 µm isotropic resolutions. Up-to-date versions of these archives can be reproduced via a script collection written and released for the purpose of this publication [14].

For evaluation, `dsurqec` and `ldsurqec` template variations (same data matrix, matched to the orientation and size requirements of the fig. 1d and fig. 1c workflow functions, respectively) are referred to as the “Generic” template. Analogously, the `ambmc` and `lambmc` template variations are referred to as the “Legacy” template.

### Qualitative Operator Inspection

We complement automated whole-dataset evaluation metrics with convenience functions to ease and improve qualitative inspection. These functions produce paginated slice-by-slice views of registered data, highlighting two different assessments. The first view mode highlights single-session registration quality, plotting the registered data as a map, and the target atlas as a contour (figs. S1 and S2). The second highlights multi-session registration coherence, plotting the target template as a map, and colored individual session contours (fig. S3).

## Evaluation

A major challenge of registration QC is that a perfect mapping from the image to the template is undefined. Similarity metrics are ill-suited for QC, as they are used internally by registration functions, whose mode of operation is based on optimizing them. Moreover, similarity metrics are not independent, so similarity score optimization issues cannot be circumvented by selecting a subset of metrics and performing QC via the remainder. To address this challenge, we developed four alternative evaluation metrics: volume conservation, smoothness conservation, functional analysis, and variance analysis. In order to mitigate possible differences arising from template features, we use these metrics for multi-factorial analyses — including both a template and a workflow factor. In order to evaluate robustness with respect to data contrast, we additionally present a multi-factorial break-down of metric variability given contrast combinations of the functional data (BOLD and CBV) and associated structural data (T1-weighted FLASH and T2-weighted RARE).

### Volume Conservation

Volume conservation is based on the assumption that brain volume should remain roughly constant after preprocessing. Assuming size consistency between data and template, a volume increase may indicate that the brain was stretched to fill template space not covered by the scan, while a volume decrease might indicate that non-brain voxels were introduced into the template brain space. For this analysis we compute a Volume Conservation Factor (VCF), whereby volume conservation is optimal for VCF = 1.

As seen in fig. 3a (left panel), we note that VCF is sensitive to the workflow (*F*_1,388_ = 182.8, *p* = 2.12 *×* 10^*−*34^), the template (*F*_1,388_ = 569.3, *p* = 4.28 *×* 10^*−*78^), but not the interaction thereof (*F*_1,388_ = 0.56, *p* = 0.45). The performance of the Generic SAMRI workflow in conjunction with the Generic template is significantly different from that of the Legacy workflow in conjunction with the Legacy template, yielding a two-tailed p-value of 1.1 *×*10^*−*20^. Moreover, the root mean squared error ratio strongly favours the Generic workflow (RMSE_L_*/*RMSE_G_ ≃2.7).

**Figure 3:**
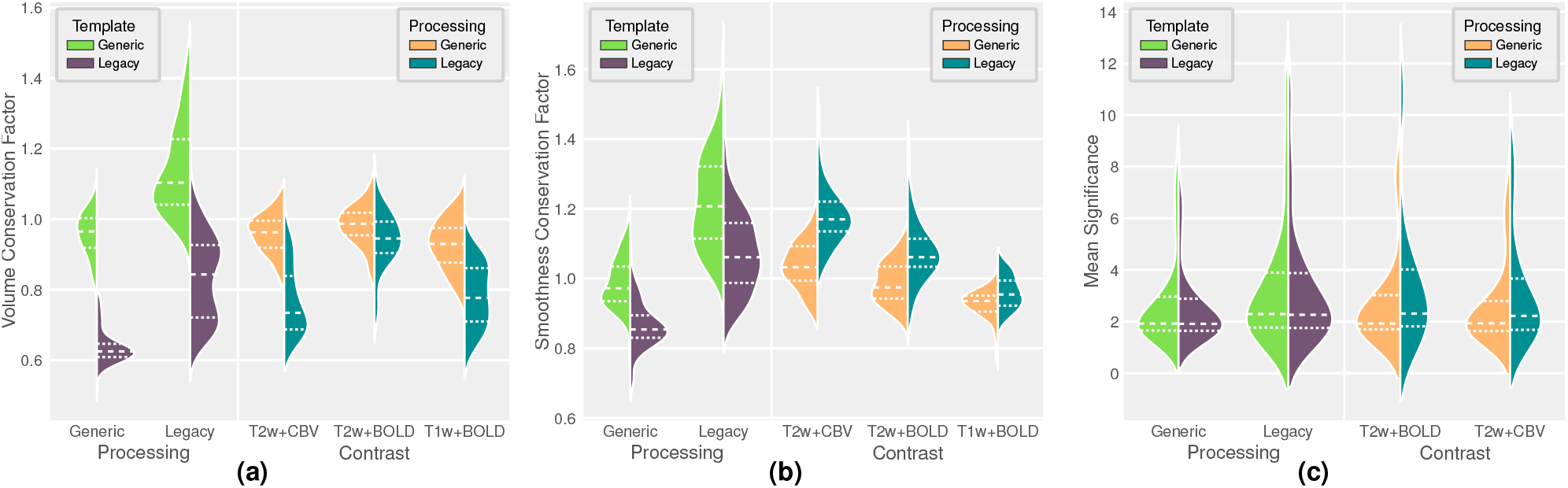
The SAMRI Generic workflow and template reliably conserve volume and smoothness — unlike the Legacy workflow and template. Plots of three target metrics, with coloured patch width distribution densities, solid line means, and dashed line inner quartiles. Comparisons across processing and templates include all contrast combinations, comparisons across processing and contrasts include only matching template-workflow combinations.

Descriptively, we observe that the Legacy level of the template variable introduces a notable volume loss (VCF of −0.32, 95%CI: −0.34 to −0.30), while the Legacy level of the workflow variable introduces a volume gain (VCF of 0.18, 95%CI: 0.16 to 0.20). Further, we note that there is a very strong variance increase in all applications of the Legacy workflow (5.9-fold given the Legacy template, and 3.9-fold given the Generic template).

With respect to data break-up by contrast (fig. 3a, right panel), we see significant effects for the contrast (*F*_2,190_ = 4.026, *p* = 0.019), and the processing-contrast interaction (*F*_2,190_ = 17.05, *p* = 1.55 *×* 10^*−*7^) factors. Multiple-comparison corrected post-hoc t-tests reveal that the processing factor remains significant in interaction with the T2w+BOLD (*p* = 0.025), T2w+CBV (*p* = 1*×*10^*−*19^), and T1w+BOLD (*p* = 9*×*10^*−*12^) levels of the contrast factor.

### Smoothness Conservation

A further aspect of preprocessing quality is the resulting image smoothness. Although controlled smoothing is a valuable tool used to increase the signal-to-noise ratio (SNR), uncontrolled smoothing limits operator discretion in the trade-off between SNR and feature granularity, leading to undocumented loss of spatial resolution and thus inferior anatomical alignment [15]. To ascertain the extent of uncontrolled smoothing, we compute a Smoothness Conservation Factor (SCF), expressing the ratio between the smoothness of the preprocessed images and the smoothness of the original images.

With respect to the data shown in fig. 3b (right panel), we note that SCF is sensitive to the template (*F*_1,388_ = 69.48, *p* = 1.35 *×*10^*−*15^), the workflow (*F*_1,388_ = 260.2, *p* = 3.64 *×* 10^*−*45^), but not the interaction of the factors (*F*_1,388_ = 1.365, *p* = 0.24). The performance of the Generic SAMRI workflow in conjunction with the Generic template is significantly different from that of the Legacy workflow in conjunction with the Legacy template, yielding a two-tailed p-value of 2.7 *×* 10^*−*19^. In this comparison, the root mean squared error ratio favours the Generic workflow (RMSE_L_*/*RMSE_G_ ≃1.9).

Descriptively, we observe that the Legacy level of the template variable introduces a smoothness reduction (SCF of −0.12, 95%CI: −0.14 to −0.10), while the Legacy level of the workflow variable introduces a smoothness gain (SCF of 0.24, 95%CI: 0.22 to 0.25). Further, we note that there is a strong variance increase for the Legacy workflow (3.08-fold given the Legacy template and 3.46-fold given the Generic template).

Given the break-up by contrast shown in fig. 3b (right panel), we see significant effects for the contrast factor (*F*_2,190_ = 21.55, *p* = 3.68*×* 10^*−*9^) and the processing-contrast interaction (*F*_2,190_ = 9.576, *p* = 0.00011). Multiple-comparison corrected post-hoc t-tests reveal that the processing factor remains significant in interaction with the T2w+BOLD (*p* = 1 *×*10^*−*6^), T2w+CBV (*p* = 2 *×* 10^*−*14^), and T1w+BOLD (*p* = 0.046) levels of the contrast factor.

### Functional Analysis

Functional analysis is a frequently used avenue for pre-processing QC, and has some viability as it is a feature not weighted by the registration process. However, the method is computationally intensive and only applicable to stimulus-evoked functional data. Additionally, functional analysis significance scores are sensitive to data smoothness [16], and thus an increased score on account of uncontrolled smoothing can be expected. For this analysis we compute the Mean Significance (MS), expressing the significance detected across all voxels of a scan, for the subset of contrasts which feature stimulus-evoked activity (T2w+CBV and T2w+BOLD).

As seen in fig. 3c (left panel), MS is sensitive to the workflow (*F*_1,268_ = 4.001, *p* = 0.046), but not to the template (*F*_1,268_ = 0.054, *p* = 0.82), or the interaction of the factors (*F*_1,268_ = 1.01*×*10^*−*6^, *p* = 1).

The performance of the SAMRI Generic workflow (with the Generic template) differs significantly from that of the Legacy workflow (with the Legacy template) in terms of MS, yielding a two-tailed p-value of 2.8*×*10^*−*9^.

Descriptively, we observe that the Legacy level of the workflow variable introduces a notable significance increase (MS of 0.70, 95%CI: 0.53 to 0.87), while the Legacy level of the template variable (MS of −0.08, 95%CI: −0.25 to 0.09), and the interaction of the Legacy template and Legacy workflow (MS of 0.00, 95%CI: −0.24 to 0.24) introduce no significance change. Furthermore, we again note a variance increase in all applications of the Legacy workflow (2.4-fold given the Legacy template, and 2.3-fold given the Generic template).

With respect to data break-up by contrast (fig. 3c, right panel), we see no notable main effect for the contrast variable (*F*_1,132_ = 0.03, *p* = 0.86), nor the contrast-template interaction (*F*_1,132_ = 0.091, *p* = 0.76).

Functional analysis effects can further be inspected by visualizing population-level maps. Second-level t-statistic maps depicting the T2w+CBV and T2w+BOLD omnibus signal (common to all subjects and sessions) provide a succinct overview capturing signal amplitude, directionality, and coverage (fig. 4). We note that the Legacy workflow induces coverage overflow (figs. 4d,4g and 4h), while the Legacy template causes acquisition slice misalignment (figs. 4b, 4d, 4f and 4h). Positive activation of the Raphe system, most clearly disambiguated from the surrounding tissue in the T2w+BOLD contrast, is notably displaced very far caudally by the joint effects of the Legacy workflow and the Legacy template (fig. 4h). Processing with the Generic template and workflow (figs. 4a and 4e) successfully mitigates all of these issues.

**Figure 4:**
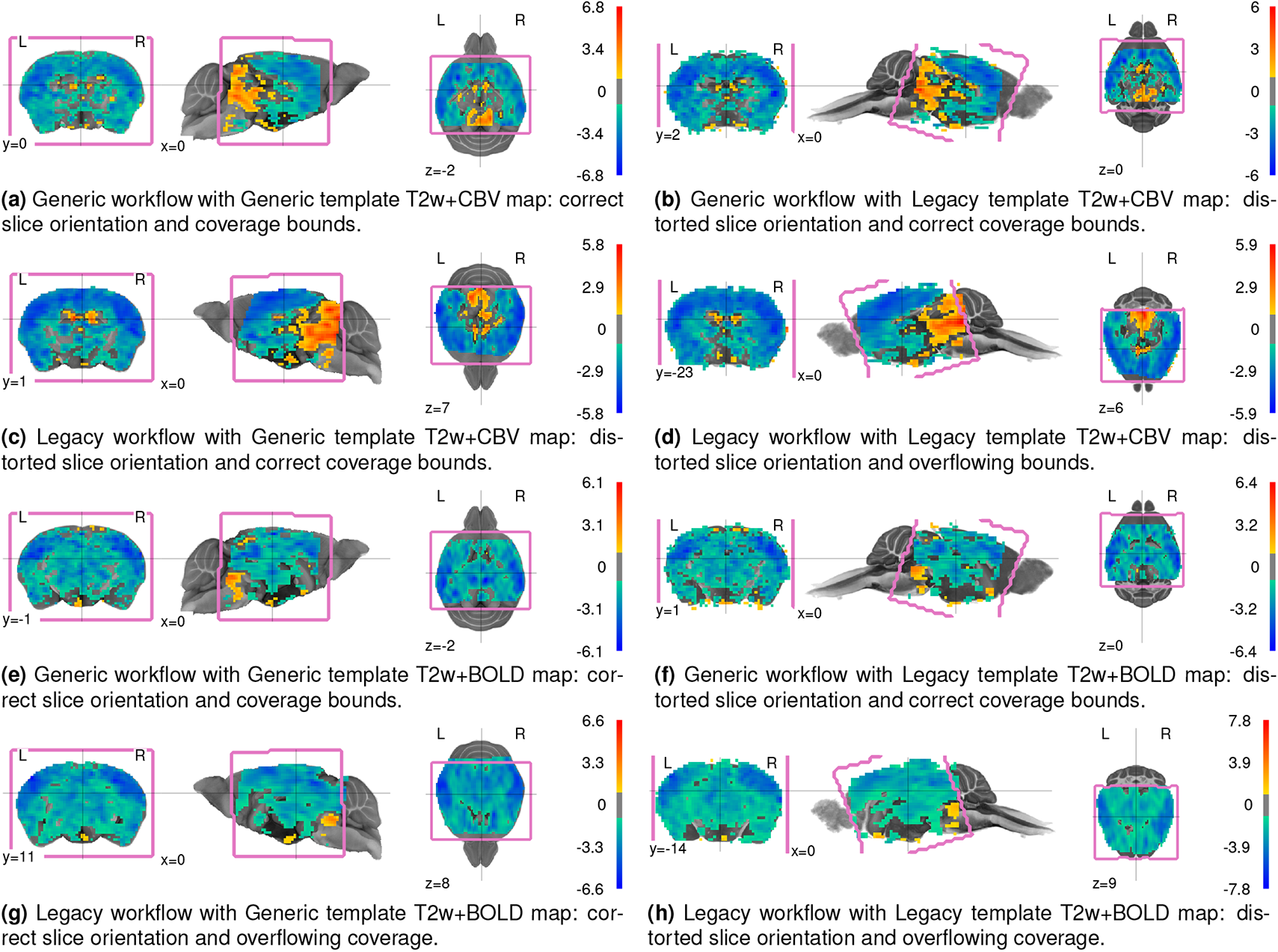
Generic processing eliminates map overflow into adjacent anatomical regions, and Generic template usage reduces slice orientation distortions. As seen in multiplanar depictions of second-level omnibus statistic maps, thresholded at |t| *≥* 1. The acquisition area is bracketed in pink, and comparison with statistic coverage should account for the latter being more constrained, as the omnibus statistic contrast is only defined for voxels captured in *every* evaluated scan.

### Variance Analysis

An additional assessment of preprocessing quality focuses on the robustness to longitudinal data variability, and whether this is attained without overfitting (i.e. compromising physiologically meaningful variability). The core assumption of this analysis is that registration should be less susceptible to experiment repetition and aging over an 8-week period in adult mice, than to inter-individual differences. Consequently, similarity scores of preprocessed scans with respect to the target template should show more variation across subject variable levels than session variable levels. This comparison can be performed using a type 3 ANOVA, modelling both the subject and the session variables. For this assessment we select three metrics: Neighborhood Cross Correlation (CC, sensitive to localized correlation), Global Correlation (GC, sensitive to whole-image correlation), and Mutual Information (MI, sensitive to whole-image information similarity).

Figure 5 renders similarity scores for the SAMRI Generic and Legacy workflows (considering only the matching workflow-template combinations). The Legacy workflow produces results with predominantly higher F-statistics for the session than the subject variable: CC (subject: *F*_10,19_ = 8.076, *p* = 0.00014, session: *F*_4,19_ = 4.659, *p* = 0.0086), GC (subject: *F*_10,19_ = 1.536, *p* = 0.22, session: *F*_4,19_ = 4.444, *p* = 0.011), and MI (subject: *F*_10,19_ = 1.597, *p* = 0.2, session: *F*_4,19_ = 4.014, *p* = 0.016).

**Figure 5:**
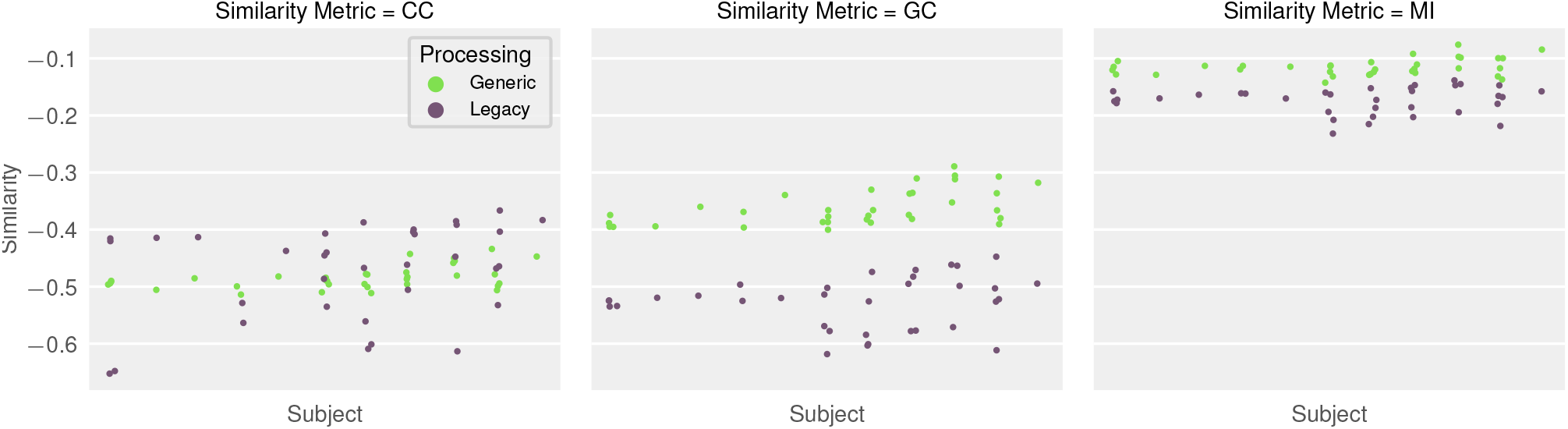
The SAMRI Generic workflow conserves subject-wise variability and minimizes trial-to-trial variability compared to the Legacy workflow. Swarmplots illustrate similarity metric scores between preprocessed images and the corresponding workflow template, plotted across subjects (x-axis bins) and sessions (individual points in each x-axis bin), for the CBV contrast.

The Generic SAMRI workflow shows a reversing trend, with overall higher F-statistic scores for the subject variable than for the session variable: CC (subject: *F*_10,19_ = 24.01, *p* = 3.65 *×*10^*−*8^, session: *F*_4,19_ = 3.456, *p* = 0.028), GC (subject: *F*_10,19_ = 5.687, *p* = 0.0012, session: *F*_4,19_ = 1.786, *p* = 0.17), and MI (subject: *F*_10,19_ = 2.258, *p* = 0.075, session: *F*_4,19_ = 2.787, *p* = 0.056).

## Discussion

The workflow and template design presented herein represent a significant methodological improvement in pre-clinical MRI, with both prospective and retroactive applicability, in conjunction with BIDS data or vendor data and vendor-to-BIDS [17] conversion workflows. The workflow offers significant advantages by reducing coverage overestimation, uncontrolled smoothing, and by guaranteeing session-to-session consistency. This is most prominently highlighted by Volume Conservation (fig. 3a), Smoothness Conservation (fig. 3b), and Variance Analysis (fig. 5), where the combined usage of the SAMRI Generic workflow and template outperforms all other combinations of the multi-factorial analysis. Increased region assignment validity is also revealed in a qualitative examination of higher-level functional maps (fig. 4), where only the Generic workflow and template combination provides accurate coverage of the sampled volume for stimulus-evoked fMRI data.

These benefits are robust with respect to tested contrast combinations (fig. 3), with the Generic workflow-template combination being less or equally susceptible to the contrast variable compared to the Legacy workflow-template combination. The performance of the Generic workflow is more consistent across all metrics, as demonstrated by notable standard deviation reductions for VCS, SCF, and MS. Variance analysis reveals that the Generic workflow is more robust than the Legacy workflow to session-specific preparation and positioning, as well as animal age progression over a period of 8 weeks, while better conserving subject-specific features.

Benchmarking data, including animals with optogenetic implants, documents workflow robustness with respect to both magnetic susceptibility artefacts (arising from scar tissue around the dental cement attachment), as well as volumetric distortion resulting from brain lesion and implant presence. Further, this last feature reveals reliable multi-session implant position conservation for the Generic (fig. S3) but not the Legacy (fig. S4) workflow. Robustness to further sources of data variability is substantiated by SNR heterogeneity in the benchmarking data collection, which is acquired not only with different scan protocols and with or without a contrast agent, but also with different coils (in-house T/R surface coil for T2w+BOLD and T2w+CBV, and cryogenic coil for T1w+BOLD). In fact, the surface coil protocol constitutes a challenging and unfavourable setting for registration due to its intrinsic *B*_1_ gradient. In spite of this, registration remains robust. Lastly, the registration pipeline contains an aggressive bias field correction step — rendering further homogeneity variations (in the spatial resolution range obtained by using different field strengths or resonator coils) mathematically equivalent to SNR variations.

Closer model inspection reveals that in addition to the processing factor, the template factor also drives variability. The Legacy template induces both volume and smoothness decrease below raw data values (figs. 3a and 3b). This clearly indicates a whole-volume effect, whereby a target template smaller than the recoded brain causes contraction during registration, affecting both volume and smoothness. We thus highlight the importance of an appropriate template choice, and strongly recommend usage of the Generic template on account of better scale similarity to data acquired in adult mice.

The volume conservation, smoothness conservation, and session-to-session consistency of the SAMRI Generic workflow and template combination are further comple-mented by numerous design benefits (figs. 1 and 2). These include increased transparency and parameterization (easing inspection and further customization), veracity of resulting data headers, and spatial coordinates more meaningful for surgical interventions.

### Quality Control

A major contribution of this work is the implementation of multiple metrics providing simple and thorough QC for registration performance (VCF, SCF, MS, and Variance Analysis); and the release of a data collection [18,19] suitable for multifactorial benchmarking.

The VCF and SCF provide good quantitative estimates of distortion prevalence. The analysis comparing subject-wise and session-wise variance is an elegant avenue for ascertaining how much a registration workflow is potentially overfitting. These metrics are relevant to both preclinical and clinical MRI workflow improvements.

Global statistical power is not a reliable metric for registration optimization. Regrettably, however, it may be the most prevalently used if results are only inspected at a higher level — and could bias analysis. This is exemplified by the positive main effect of the Legacy workflow seen in fig. 3c. In this particular case, optimizing for statistical power alone would give a misleading indication, as becomes evident when all metrics are inspected.

We suggest that a VCF, SCF and Variance based comparison, coupled with visual inspection of a small number of omnibus statistic maps is a feasible and sufficient tool for benchmarking workflows. We recommend reuse of the presented data for workflow benchmarking, as they include (a) multiple sources of variation (contrast, session, subjects), (b) functional activity with broad coverage but spatially distinct features, and (c) significant distortions due to implant properties — which are appropriate for testing workflow robustness. In addition to the workflow code [10], we openly release the re-executable source code [20] for all statistics and figures in this document. Thus both the novel method and the benchmarking process are fully transparent and reusable with further data.

### Conclusion

We present a novel registration workflow, “SAMRI Generic”, which offers several advantages compared to ad hoc approaches commonly used for small animal MRI. In-depth multivariate comparison with a thoroughly documented Legacy pipeline revealed superior performance of the SAMRI Generic workflow in terms of volume and smoothness conservation, as well as variance structure across subjects and sessions. The metrics introduced for registration QC are not restricted to the processing of small animal fMRI data, and can be readily expanded to other brain imaging applications. The optimized registration parameters of the SAMRI Generic Workflow are easily accessible in the source code and transferable to any other workflows making use of the ANTs package. Overall, we believe that the SAMRI Generic workflow should facilitate and harmonize processing of mouse brain imaging data across studies and centers.

## Methods

### Data

For the quality control of the workflows, a dataset with an effective size of 164 scans is used. Scans were acquired using a Bruker PharmaScan system (7 T, 16 cm bore), and variable coils. Experimental procedures were approved by the Veterinary Office of the Canton of Zurich in accordance with relevant regulations.

#### T2w+CBV and T2w+BOLD

Data was acquired from 11 adult animals (6 males and 5 females, 207 days to 261 days old at study onset), with each animal scanned on up to 5 sessions (at 14 day intervals). Each session contains an anatomical scan and two functional scans — with Blood-Oxygen Level Dependent (BOLD) [21] and Cerebral Blood Volume (CBV) [22] contrast, respectively (for a total of 68 functional scans).

Anatomical scans were acquired via a TurboRARE sequence, with a RARE factor of 8, an echo time (TE) of 21 ms, an inter-echo spacing of 7 ms, and a repetition time (TR) of 2500 ms, sampled sagittally at Δx(*ν*) = 166.7 µm, horizontally at Δy(*ϕ*) = 75 µm, and coronally at Δz(t) = 650 µm (slice thickness of 500 µm). The field of view covered 20 *×* 9 mm^2^ and was sampled via a 120 *×* 120 matrix. A total of 14 slices were acquired.

The functional BOLD and CBV scans were acquired using a gradient-echo Echo Planar Imaging (ge-EPI) sequence with a flip angle of 60° and with TR*/*TE = 1000 ms*/*15 ms and TR*/*TE = 1000 ms*/*5.5 ms, respectively. Data were sampled at Δx(*ν*) = 312.5 µm, Δy(*ϕ*) = 281.25 µm, and Δz(t) = 650 µm (slice thickness of 500 µm). The field of view covered 20*×* 9 mm^2^ and was sampled via a 64 *×*32 matrix. A total of 14 slices were acquired.

Animals were fitted with an optic fiber implant (l = 3.2 mm d = 400 µm) targeting the Dorsal Raphe (DR) nucleus. The nucleus was sensitized to optical stimulation by transgenic expression of Cre recombinase under the ePet promoter [23] and rAAV delivery of a plasmid for Cre-conditional Channelrhodopsin and YFP expression — pAAV-EF1a-double floxed-hChR2(H134R)EYFP-WPRE-HGHpA, reposited by Karl Deisseroth (Addgene plasmid #20298). Scans were acquired using an inhouse T/R surface coil, engineered to permit optic fiber protrusion.

The DR was stimulated via an Omicron LuxX 488-60 laser (488 nm) tuned to 30 mW at contact with the fiber implant, according to the protocol listed in table S1. Stimulation was delivered and reports recorded via the COSplay system [24].

#### T1w+BOLD

Data was acquired from 10 adult male animals (34 days old at study onset), with each animal scanned on 3 sessions (repeated at 58 days postnatal and 112 days postnatal) [25]. Each session contains an anatomical scan and a functional BOLD contrast scan (for a total of 30 functional scans).

Anatomical scans were acquired via a Fast low-angle shot (FLASH) sequence with a RARE factor of 1, a flip angle of 30°, and TR*/*TE = 522 ms*/*3.51 ms. Data were sampled at Δx(*ν*) = 52 µm, Δy(*ϕ*) = 26 µm, Δz(t) = 500 µm (slice thickness of 500 µm). The field of view covered 20 *×* 10 mm^2^ and was sampled via a 384 *×* 384 matrix. A total of 20 slices were acquired.

The functional BOLD scans were acquired using a gradient-echo Echo Planar Imaging (ge-EPI) sequence with a flip angle of 60° and TR*/*TE = 1000 ms*/*15 ms. Data were sampled at Δx(*ν*) = 222 µm, Δy(*ϕ*) = 200 µm, and Δz(t) = 500 µm (slice thickness of 500 µm). The field of view covered 20 *×* 10 mm^2^ and was sampled via a 90 *×* 50 matrix. A total of 20 slices were acquired.

### Metrics

For the VCF we use the 66^th^ intensity percentile of the raw scan as a signal threshold. The arbitrary unit equivalent of this threshold is recorded per-scan and applied to all preprocessing results for that particular scan — eq. (1), where *v* is the voxel volume in the original space, *v′* the voxel volume in the transformed space, *n* the number of voxels in the original space, *m* the number of voxels in the transformed space, *s* a voxel value sampled from the vector *S* containing all values in the original data, and *s′* a voxel value sampled from the transformed data.

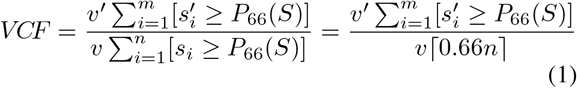

The SCF metric measures the pre-and post-processing smoothness ratio, defined as the full-width at half-maximum (FWHM) of the signal amplitude spatial auto-correlation function (ACF [26]). As fMRI data have non-Gaussian spatial ACF, we use AFNI [27] to fit the following function — eq. (2), where *r* is the distance of two amplitude distribution samples, *a* is the relative weight of the Gaussian, *b* is the width of the Gaussian, and *c* the decay of the mono-exponential term [28].

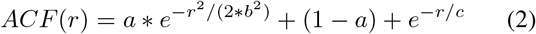

Statistical power as per the MS metric is represented by the negative logarithm of first-level p-values. This produces voxelwise statistical estimates for the probability that a time course could — by chance alone — be at least as well correlated with the stimulation regressor as the voxel time course measured. We compute the per-scan average of these values as seen in eq. (3), where *n* represents the number of statistical estimates in the scan, and *p* is a p-value.

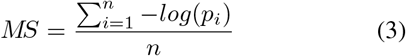

Two statistical models, including main effects and interaction terms, were evaluated for each metric. The first details workflow and template effects, and the second details contrast effects. Multiple comparison correction was performed via the Benjamini-Hochberg method, fixing the false discovery rate (FDR) *α* at 0.05.

### Software

The workflow functions leverage the Nipype [29], FSL [1] and ANTs [2] packages. Software management relevant for verifying the reproducibility of the software environment and analysis results was performed via neuroscience package install instructions for the Gentoo Linux distribution [30]. For Quality Control we distribute as part of this article workflows using the SciPy [31], pandas [32], Seaborn [33], and Statsmodels [34] packages.

## Reproducibility and Resource Availability

All data is freely available [18,19], and automatically accessible via the Gentoo Linux package manager. All code relevant for reproducing this document is freely available [20], and compliant with the RepSeP specification [35] to allow article reproduction based solely on the raw data and top-level code of the article.

## Supplementary Materials

**Table S1:**
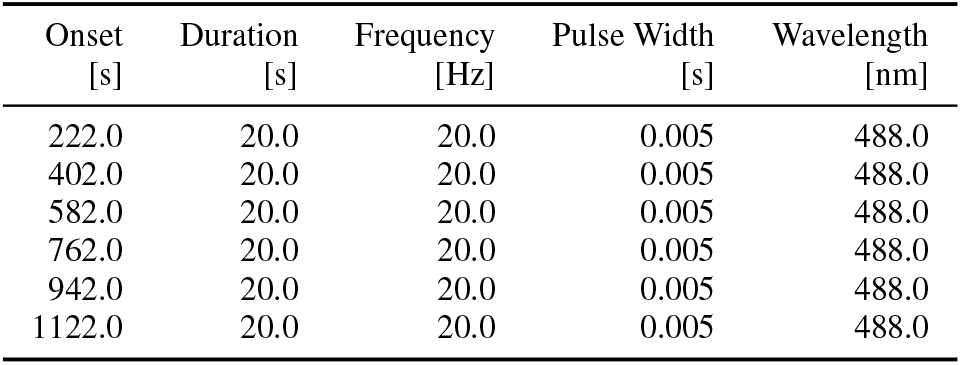
Stimulation protocol, as delivered during functional scans. Stimulus event spacing and parameters are constant across scans, but the exact onset time is variable in the 10 s magnitude range due to scanner adjustment time variability.

| Onset<br>[s] | Duration<br>[s] | Frequency<br>[Hz] | Pulse Width<br>[s] | Wavelength<br>[nm] |
| --- | --- | --- | --- | --- |
| 222.0 | 20.0 | 20.0 | 0.005 | 488.0 |
| 402.0 | 20.0 | 20.0 | 0.005 | 488.0 |
| 582.0 | 20.0 | 20.0 | 0.005 | 488.0 |
| 762.0 | 20.0 | 20.0 | 0.005 | 488.0 |
| 942.0 | 20.0 | 20.0 | 0.005 | 488.0 |
| 1122.0 | 20.0 | 20.0 | 0.005 | 488.0 |

**Figure S1:**
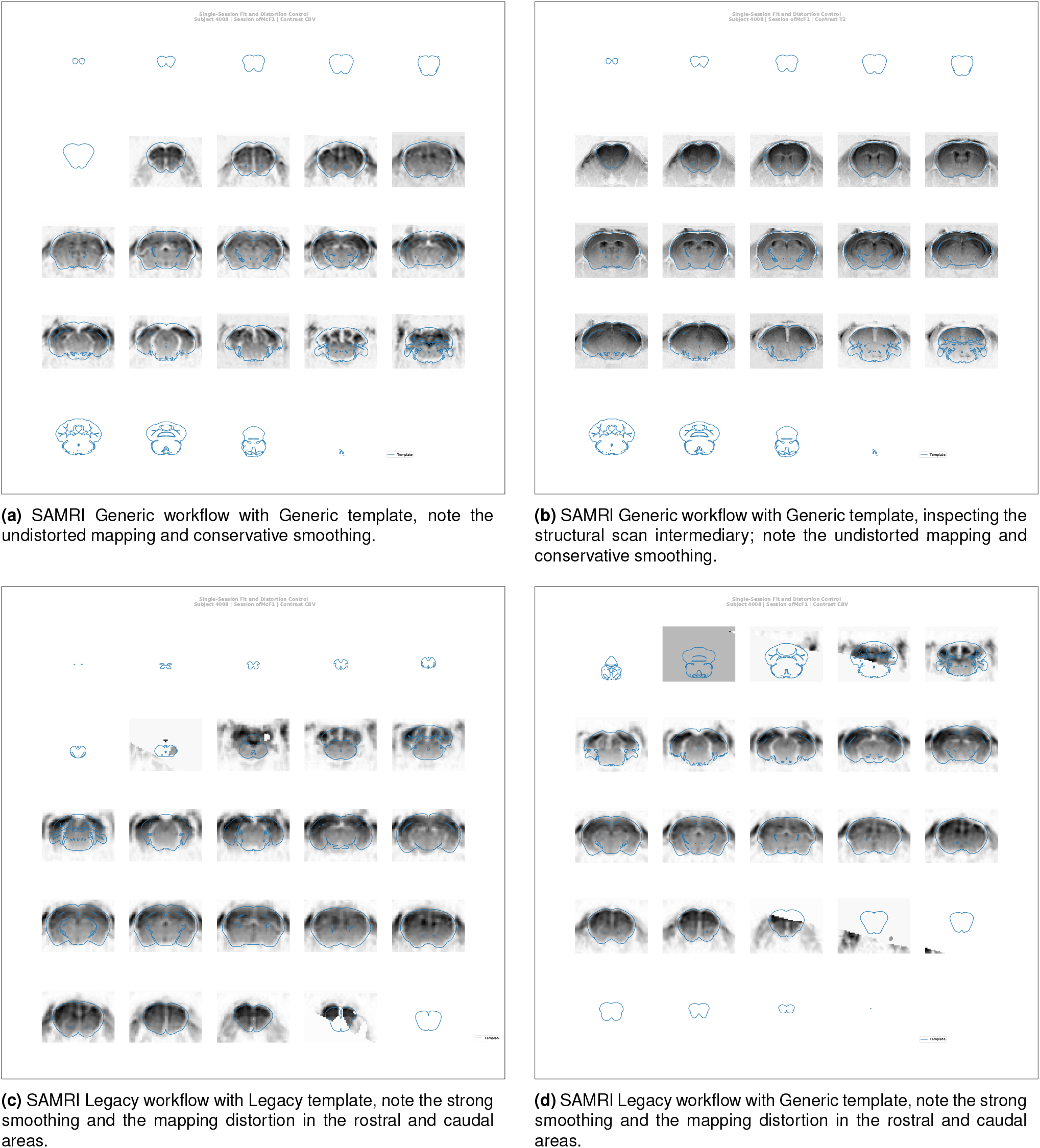
The SAMRI Generic workflow induces less smoothness, and provides more accurate coverage for CBV+T2w contrast scans. Depicted are automatically created operator overview graphics, which allow a slice-by-slice (spacing analogous to acquisition) inspection of the registration fit. This representation affords a particularly detailed view of the preprocessed MRI data, and highly accurate template contours.

**Figure S2:**
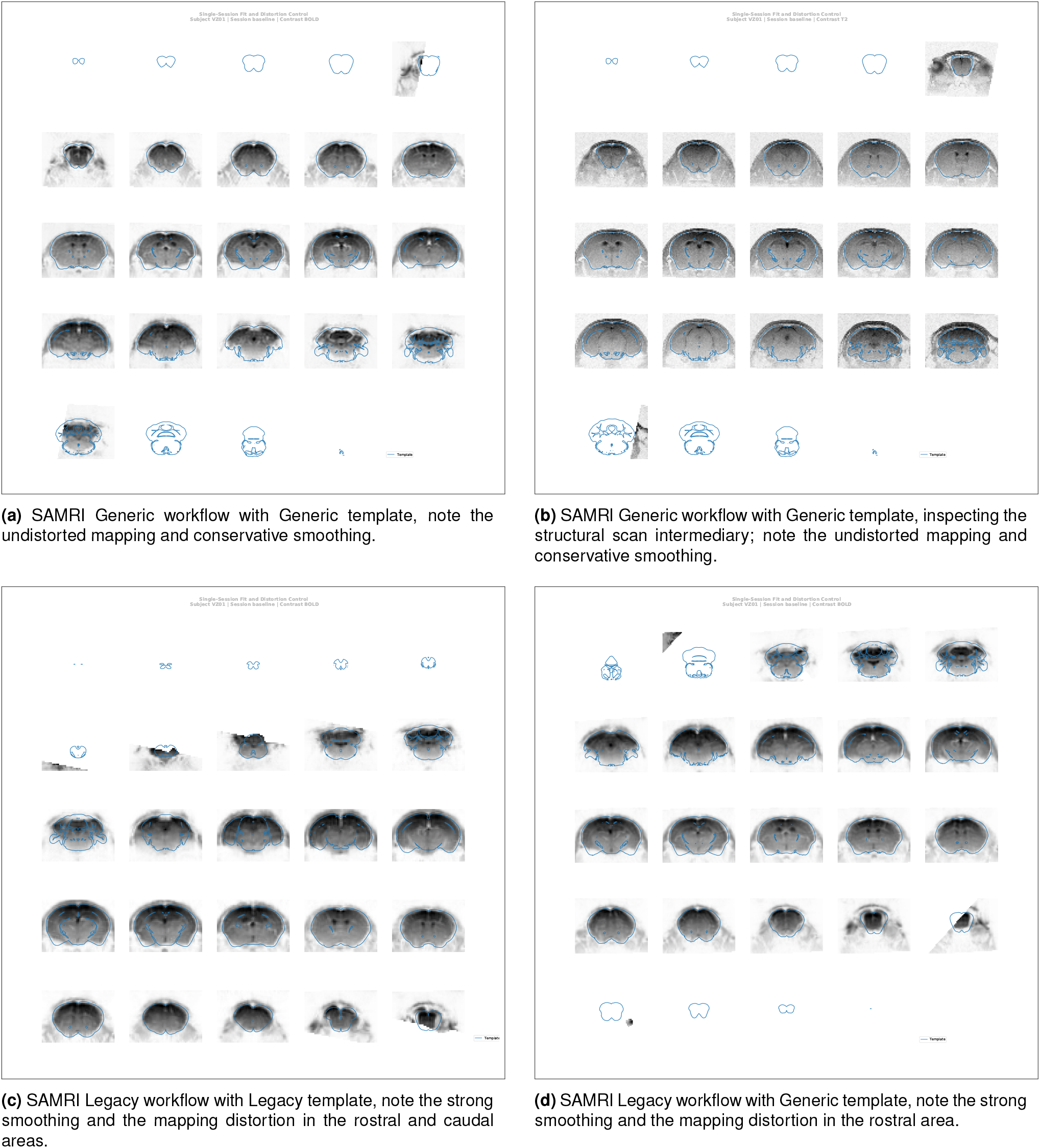
The SAMRI Generic workflow induces less smoothness, and provides more accurate coverage for BOLD+T1w contrast scans. Depicted are automatically created operator overview graphics, which allow a slice-by-slice (spacing analogous to acquisition) inspection of the registration fit. This representation affords a particularly detailed view of the preprocessed MRI data, and highly accurate template contours.

**Figure S3:**
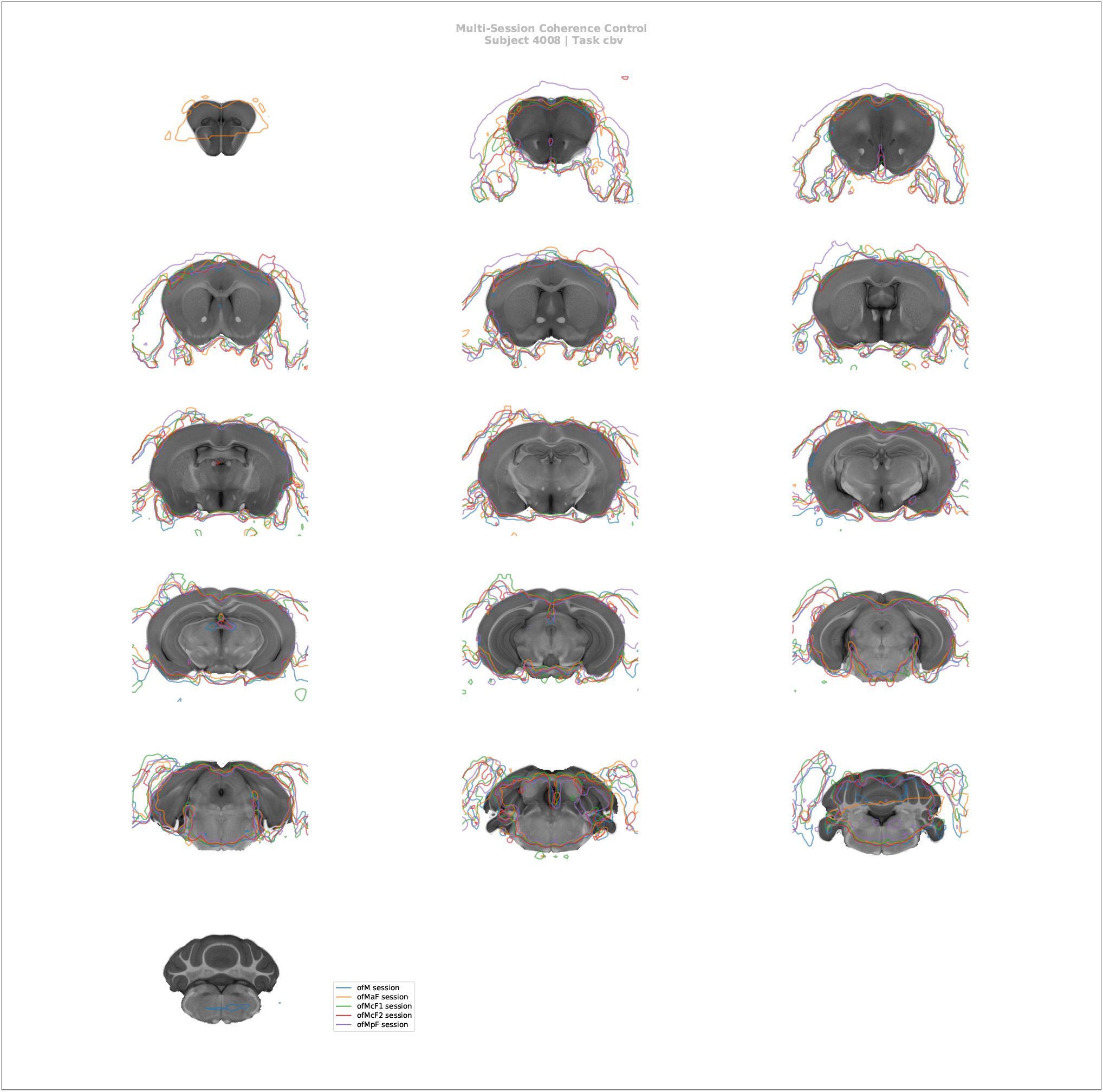
The SAMRI Generic workflow consistently maps high-salience features such as the implant site across sessions. Automatically created operator overview graphic, allowing a slice-by-slice (spacing analogous to acquisition) inspection of registration coherence. This representation permits a coarse assessment of registration consistency for multiple sessions — though at the cost of some clarity. Particularly, this visualization, allows an operator to track the position of high-amplitude fixed features across scans in order to ascertain coherence (similarly to what is automatically assessed by the Variance analysis of the session factor).

**Figure S4:**
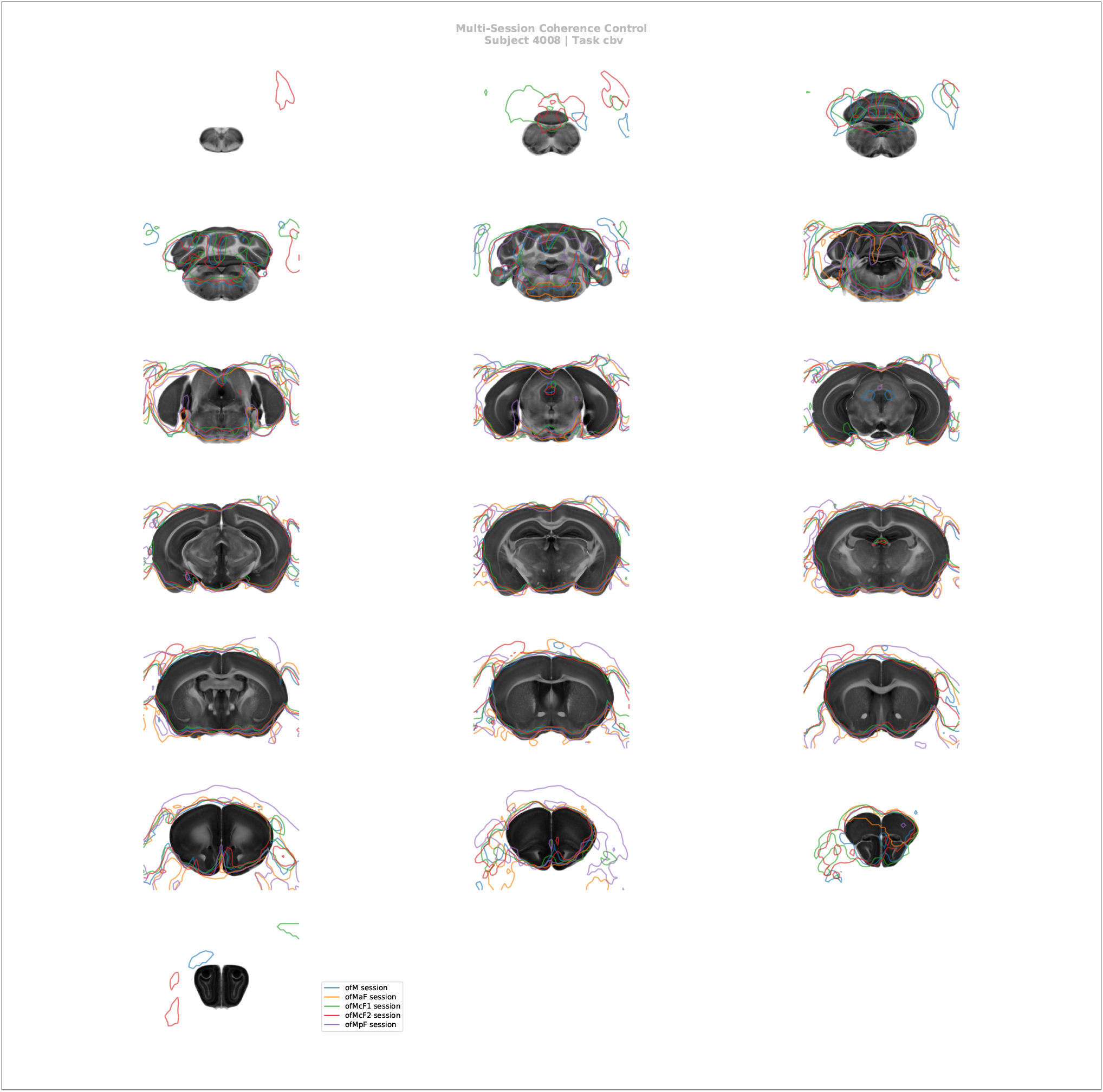
The SAMRI Legacy workflow inconsistently and incorrectly maps high-salience features such as the implant site across sessions. Automatically created operator overview graphic, allowing a slice-by-slice (spacing analogous to acquisition) inspection of registration coherence. This representation permits a coarse assessment of registration consistency for multiple sessions — though at the cost of some clarity. Particularly, this visualization, allows an operator to track the position of high-amplitude fixed features across scans in order to ascertain coherence (similarly to what is automatically assessed by the Variance analysis of the session factor).

**Figure S5:**
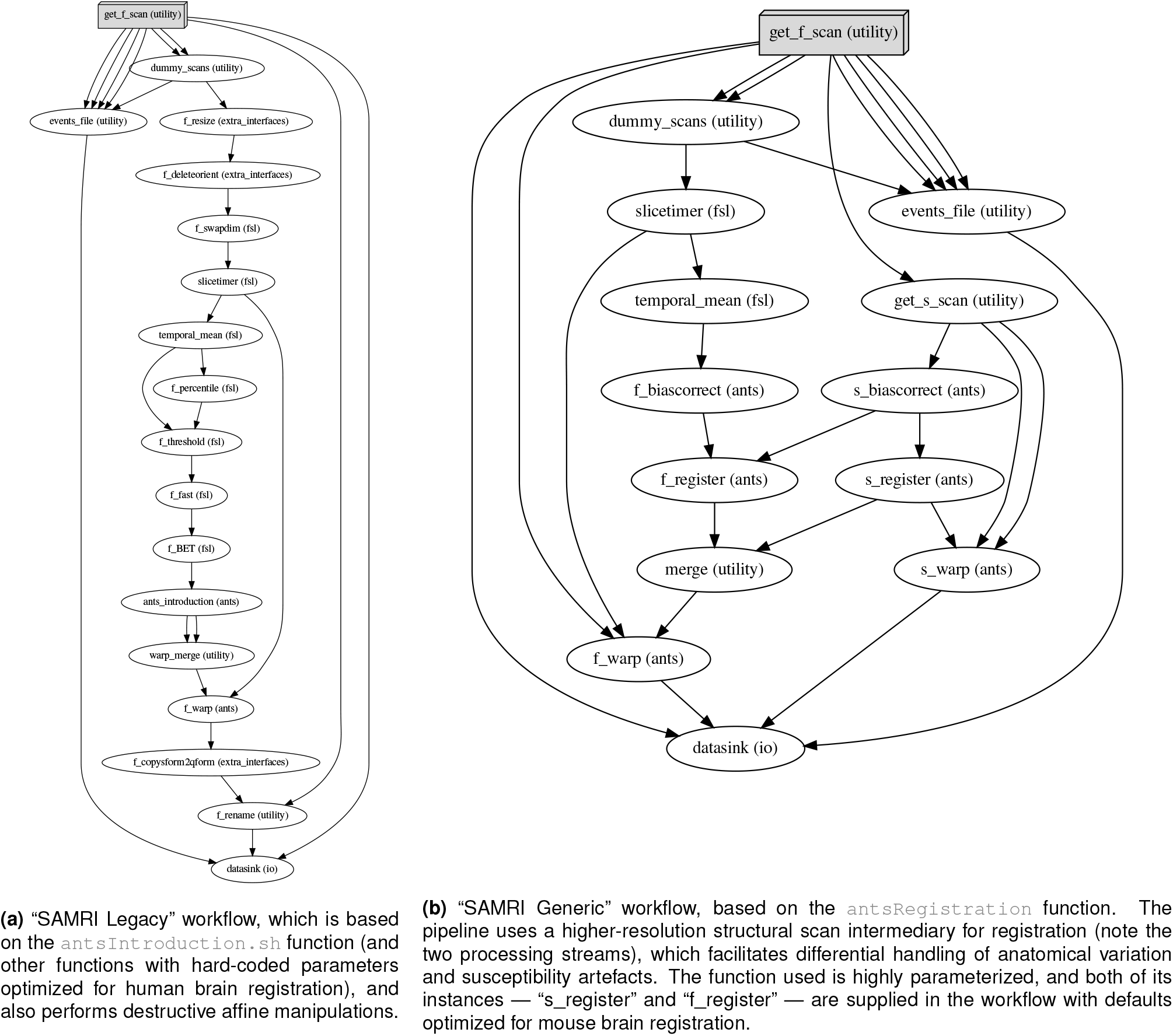
Directed acyclic graphs detailing the precise node names (as seen in the SAMRI source code) for the two alternate MRI registration workflows. The package correspondence of each processing node is appended in parentheses to the node name. The “utility” indication corresponds to nodes based on Python functions specific to the workflow, distributed alongside it, and dynamically wrapped via Nipype. The “extra_interfaces” indication corresponds to nodes using explicitly defined Nipype-style interfaces, which are specific to the workflow and distributed alongside it.

## Notes

https://bitbucket.org/TheChymera/irsabi/

https://doi.org/10.5281/zenodo.2651720

https://doi.org/10.5281/zenodo.3445428

https://doi.org/10.5281/zenodo.2545838

### Competing Interest Statement

The authors have declared no competing interest.

### Summary of Updates

Updated data collection, extended statistical modelling, reexecuted all analyses from scratch.

